# Cerebellum-Targeted Transcranial Focused Ultrasound Stimulation Modulates Hippocampus Neural Activities

**DOI:** 10.1101/2025.04.09.647932

**Authors:** Yingjian Liu, King Yi Cho, Jacky Tin Shing Hung, Caiyi Zhuo, Hei Ming Lai, Chung Tin

## Abstract

Low-intensity transcranial focused ultrasound stimulation (tFUS) has recently emerged as a promising neuromodulation technique due to its non-invasive nature, ability to penetrate deeply, and high spatial resolution. On the other hand, the cerebellum has attracted new interest as a target for neuromodulation. In this study, we investigated how cerebellum-targeted tFUS could modulate hippocampal neural activity. We found that tFUS at pulse repetition frequency (PRF) as high as 10kHz can effectively activate the cerebellum in vivo. Furthermore, PRF at 10kHz, rather than 1kHz, favored remote inhibition of hippocampus neurons. We found that the neuromodulation at hippocampus was mainly mediated by cerebellum cortex than deep cerebellar nucleus. Furthermore, by cFos expression mapping and phase-amplitude coupling analysis, we showed that the hippocampal response at different PRFs resulted from altered neurodynamic interaction rather than overall activation of engaged brain regions. Our results reveal new potential for cerebellum as a neuromodulation target for hippocampus-related functions.

## Introduction

Transcranial Focused ultrasound stimulation (tFUS) has emerged as a promising neuromodulation technique due to its non-invasiveness, deep penetration and high spatial resolution (in millimeter scale). Neuromodulation by low-intensity ultrasound represents a powerful and safe new tool for neuroscience and clinical application and has been widely investigated in small animals (Yu, Niu et al. 2021, Ramachandran, Niu et al. 2022), non-human primates (Folloni, Fouragnan et al. 2021) and humans (Legon, Sato et al. 2014). For example, it has been shown that tFUS at human primary somatosensory cortex can improve performance on sensory discrimination tasks (Legon, Sato et al. 2014). The use of low-intensity tFUS in neuromodulation has a strong potential but remains exploratory. There have been reports on the mixed effect of tFUS on the neural system. For example, Yuen et al. showed that low-intensity tFUS (500 kHz) at mouse motor cortical regions could induce tail movement, excitatory neural activity, and hemodynamic responses (Yuan, Wang et al. 2020). Conversely, Darrow et. al. showed that 3.2MHz tFUS targeting the rat thalamus reversibly suppressed somatosensory evoked potentials (Darrow, O’Brien et al. 2019).

Ultrasound stimulation can be controlled with several parameters, including fundamental frequency (FF), stimulus duration (SD), duty cycle (DC), and pulse repetition frequency (PRF). Previous studies show that PRF in the range of 1-3kHz are the most effective for stimulating neural response in the targeted area (Tufail, Yoshihiro et al. 2011). A recent study shows that excitatory and inhibitory neurons in somatosensory cortex were differentially activated by tFUS of different PRF (300 vs. 30 Hz) (Yu, Niu et al. 2021). In another study, it was shown that ultrasound at PRF=900Hz increased activity of inhibitory neurons while suppressing activity of excitatory neurons in the mouse hippocampus (Murphy, Farrell et al. 2022). There are few studies using higher PRFs but a recent work used PRF=3kHz to 10kHz to induce long-term depression (LTD) in the field excitatory postsynaptic potential (fEPSP) of rat hippocampus (Niu, Yu et al. 2022). These studies suggest that PRF can be an effective parameter to tune the neuronal response to tFUS. Yet, it remains poorly understood on how to systematically control the neuromodulation effect with appropriate PRF.

The cerebellum has traditionally been considered primarily a motor-related brain region. However, several recent studies have demonstrated that cerebellum likely contributes to a variety of cognitive functions as well, via its extensive connections with numerous cerebral regions. For example, recent studies have shown that acute excitation of the cerebellum induces altered cFos expression in hippocampus, changes in evoked local field potentials, modified hippocampal neural dynamics, and disrupted object-location processing in CA1 (Zeidler, Hoffmann et al. 2020). These findings indicate functional interactions between the cerebellum and hippocampus. The influence of cerebellum on hippocampus neural dynamics and function has important potential for epilepsy treatment. Temporal lobe epilepsy (TLE) is the most common form of epilepsy found in adults and is characterized by chronic, spontaneous seizures typically arising in the hippocampal formation (Chatzikonstantinou 2014, Streng, Tetzlaff et al. 2021). Modulation of hippocampus neural activities has been implicated in seizure suppression in many studies (Chiang, Lin et al. 2013, Davis and Gaitanis 2020). On the other hand, while not traditionally thought to be associated with epilepsy, anatomical, clinical, and electrophysiological studies suggest the cerebellum can play an important role in seizure networks (Streng and Krook-Magnuson 2021, Streng, Tetzlaff et al. 2021, Beckinghausen, Ortiz-Guzman et al. 2023, Stieve, Richner et al. 2023). Nevertheless, cerebellar electrical stimulation has produced mixed results in both human (Chkhenkeli, Sramka et al. 2004) and animal (Rucci, Giretti et al. 1968) model of epilepsy. Such inconsistency is likely due to the complexity of cell-type specific response to the stimulation, site of stimulation and parallel multi-synaptic connections from cerebellum to the site of epileptic seizure, such as the hippocampus. However, a recent study using optogenetic manipulation on the fastigial nucleus suggests that excitation, but not inhibition, of the nucleus outputs is a key determinant for the success in inhibition of TLE (Streng and Krook-Magnuson 2020, Streng, Tetzlaff et al. 2021). This suggests that with careful selection of stimulus parameters, robust seizure suppression could be possibly achieved through cerebellar stimulation.

In the work by Zeidler et. al (Zeidler, Hoffmann et al. 2020), the effect of acute cerebellum stimulation on hippocampus was investigated with optogenetic stimulation. However, the effect of tFUS neuromodulation by targeting the cerebellum has not been studied. In this study, we aimed to examine the neural response of hippocampus by stimulating the cerebellum with tFUS. In particular, we compared the effect of two different PRFs of tFUS. Few studies have used PRF up to 10kHz for neuromodulation. But we found that the cerebellum was robustly activated by the tFUS at PRF of 10kHz. We also observed that different PRF had different impacts on the neural dynamics in the cerebellum-hippocampus interaction. Together, our findings provide new key insight in the potential of tFUS for neuromodulation and broaden our understanding on the cerebellar influence on hippocampus.

## Results

### 1. Cerebellum-targeted tFUS elicited neural response in hippocampus

The *in vivo* experimental setup and waveform of the ultrasound stimulus are illustrated in Fig. 1. The ultrasound transducer (Model V301-SU, Olympus, USA) was positioned at an incidence angle of 40° over the cerebellum (Fig. 1A). A 3D printed collimator was used to focus the ultrasound wave while penetrating the skull and the brain tissue. Neural responses were recorded using a 16-channel micro-electrode array implanted in the cerebellum and/or hippocampus. As shown in Fig. 1B, pulsed ultrasound stimulus was used with fixed fundamental frequency (FF = 500 kHz), ultrasound duration (UD = 500ms) and interstimulus interval (ISI = 10sec). We used two different values of pulse repetition frequency (PRF = 1kHz or 10kHz) in our experiments. We adjusted the burst duration accordingly to maintain a constant duty cycle (DC = 50%).

**Fig. 1.**
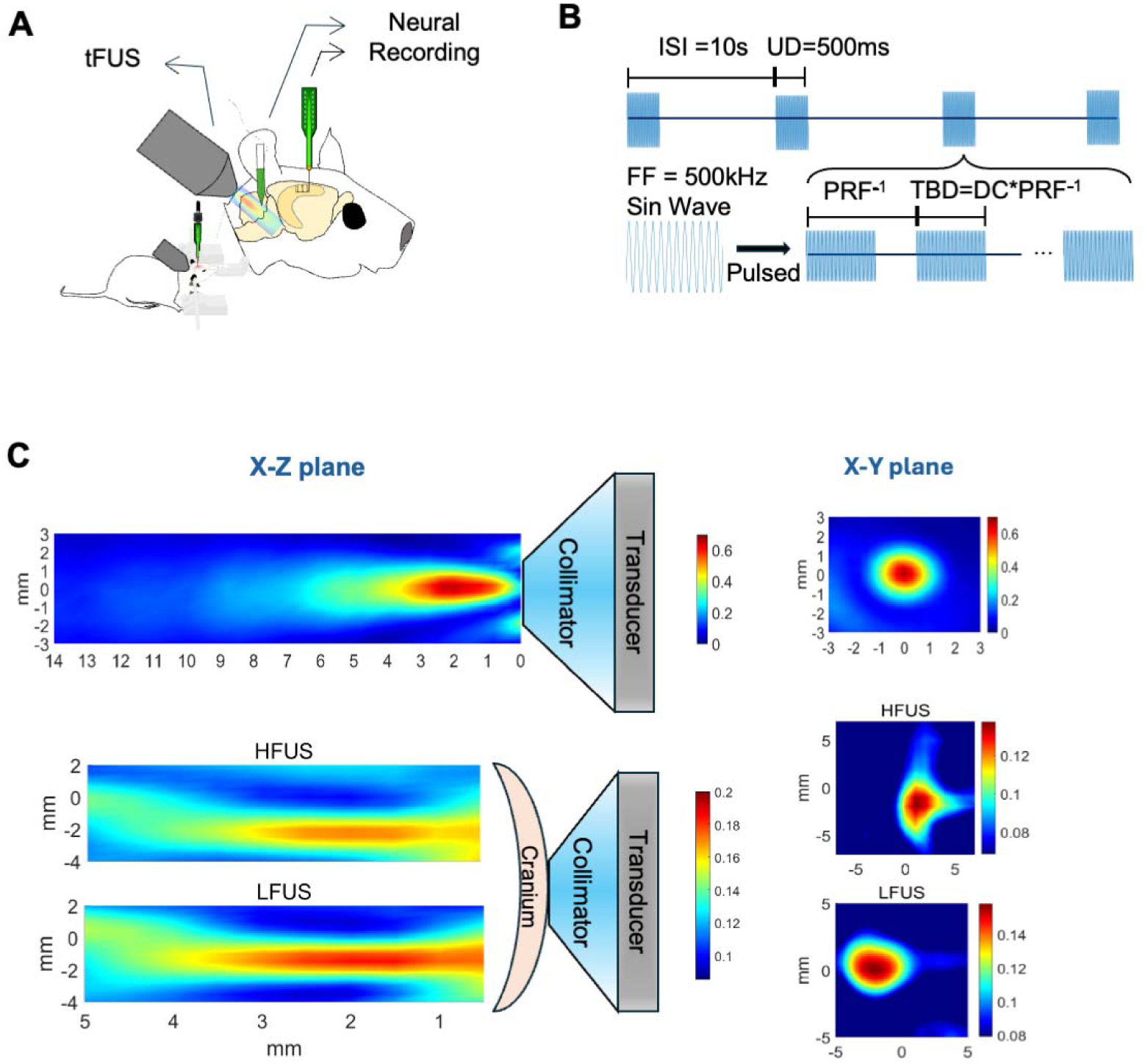
Experimental setup of ultrasound stimulation and electrophysiological recording. (A) Ultrasound stimulation targets at the cerebellum (ML: −4 mm, AP: −15 mm) at 40° incidence angle. Electrophysiological recordings were done using a 16-channel microelectrode array (MEA) in the cerebellum (ML: −4 mm, AP: −10 mm, DV: -3mm) and/or hippocampus (ML: −2.3 mm, AP: −3 mm, DV: −3 mm). (B) Key parameters of ultrasound stimulation used: fundamental frequency (FF), pulse repetitive frequency (PRF), duty cycle (DC), tone-burst duration (TBD), interstimulus interval (ISI), and ultrasound duration (UD). (C) Hydrophone-measured ultrasound field distributions in transverse (X–Y) and sagittal (X–Z) planes, with and without cranial bone, using 3D-printed collimator guidance.

Spatial characterization of the tFUS profile was mapped and reconstructed using a hydrophone-based (HCL-0200, Onda, USA) 3D ultrasound pressure scanning system (AG-2010, Onda, USA), both with and without a freshly excised rat cranium using two different PRFs (Fig. 1C). The transverse (X–Y plane) view reconstruction revealed PRF-dependent ultrasound scattering patterns through the rat cranium. Distortion of the acoustic field by the cranium was more pronounced at PRF = 10kHz. However, in both cases, the acoustic field maintained a concentric profile. The penetration depth of the acoustic wave was slightly higher with PRF = 1kHz (sagittal view, X-Z plane). The radial scattering from the penetration axis was similar under both PRFs. The spatial-peak pulse-average intensity (I_sppa_) was found to be 1.53W/cm^2^ without the cranium and 0.35W/cm^2^ with the cranium, respectively. We confirmed with H&E staining that no tissue damage was found in the brain after stimulation with these parameters (Supplementary Fig.1A). The stimulation intensity used should be below the threshold of thermal damage.

Our results show that cerebellum-targeted tFUS (Fig. 2A) in rats was able to robustly elicit neural response in hippocampus (Fig. 2B, left panels in red). Increased in firing rate of hippocampal neurons was observed shortly after tFUS at cerebellum.

**Fig. 2.**
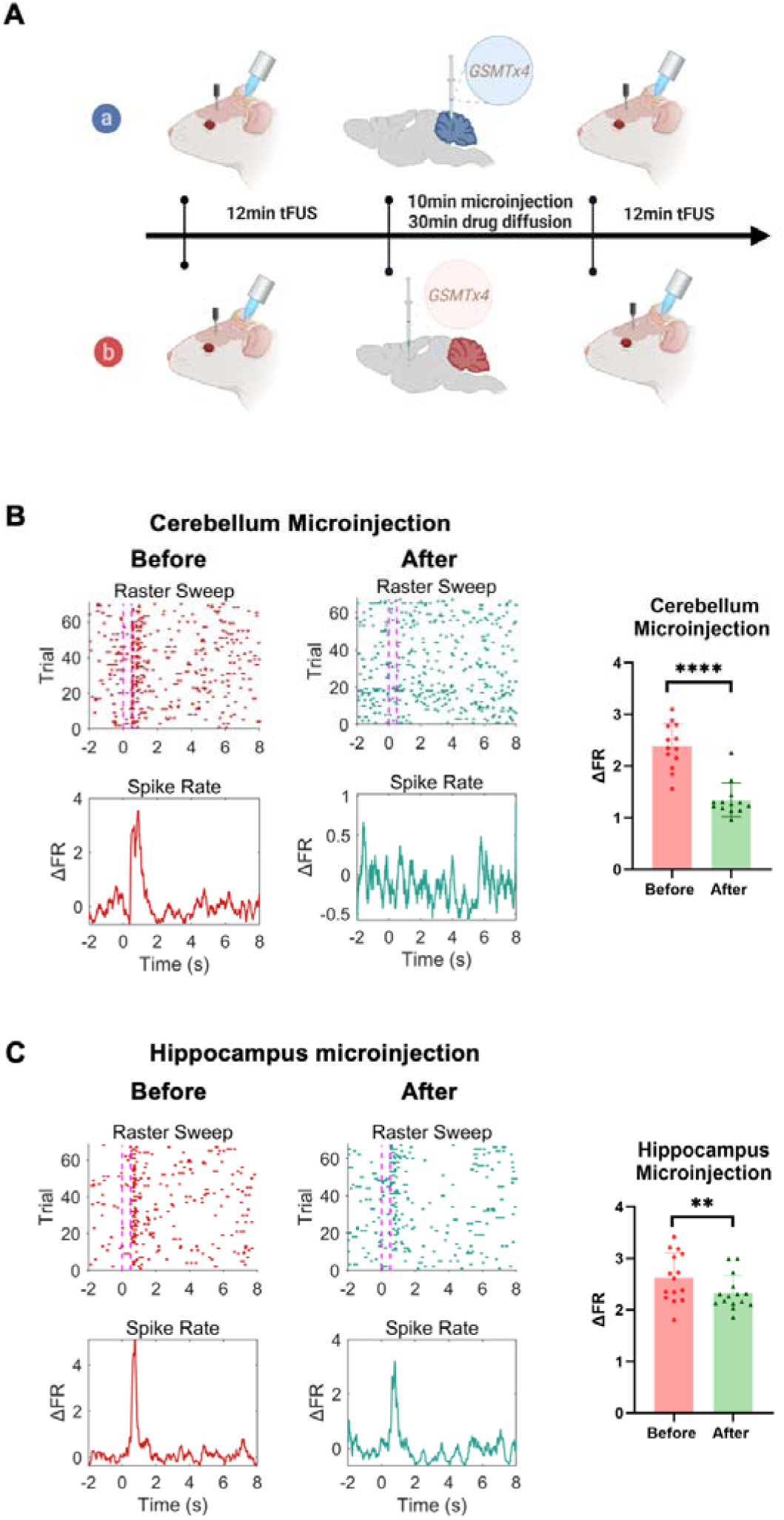
Cerebellum-targeted tFUS elicits hippocampal neural response remotely. (A) GsMTx4 (250 μmol/L, 0.5 μL/rat) was microinjected into either cerebellum (ML: −4, AP: −10, DV: −3 mm; N=3) or hippocampus (ML: −2.3, AP: −3, DV: −3 mm; N=3). (B&C) Representative neural units recorded before and after GsMTx4 microinjection in cerebellum (B) and hippocampal (C), respectively, subject to cerebellum-targeted tFUS. Left: Peristimulus raster plots, Right: Relative firing rate (ΔFR). (paired t-test, **p < 0.01, ****p < 0.0001, N = 3 rats per group, n = 15 neural units per group).

Nevertheless, previous study showed that acoustic reflection within the skull may also elicit neural response in hippocampus directly rather than via the neural circuity pathway from cerebellum. To distinguish these two possibilities, we injected GsMTx4, a selective inhibitor of cationic mechanosensitive channels including TRP and Piezo families (Bae, Sachs et al. 2011, Gnanasambandam, Ghatak et al. 2017), into the hippocampus or cerebellum, respectively (Fig. 2A). It has been shown that low-intensity tFUS primarily induced neural response *in vivo* through mechanosensitive ion channels (Dell’Italia, Sanguinetti et al. 2022). Several channels, such as Piezo (Qiu, Guo et al. 2019), TRPV (Yang, Pacia et al. 2021, Xu, Yang et al. 2023), and Mscl-G22S (Qiu, Kala et al. 2020), have been found to serve as ultrasonic amplifiers. GsMTx4 can effectively blocks ultrasound-induced neuronal responses in the central nervous system (Yoo, Mittelstein et al. 2022). Hippocampal neurons were recorded before and after GsMTx4 microinjection at the cerebellum (Fig.2B) and hippocampus (Fig.2C), respectively. Fig. 2B&C show that microinjection of GsMTx4 at cerebellum weakened the hippocampus neural response much more than microinjection at hippocampus (at cerebellum: ΔFR = 2.45 ± 0.2 to 1.34 ± 0.3, n=15, p < 0.0001; at hippocampus: ΔFR = 2.69 ± 0.7 to 2.13 ± 0.3, n=15, p < 0.01). Together, it suggests that the hippocampal neural response observed in our results should be primarily via the neural pathway from cerebellum instead of direct acoustic effect from reflection within the skull.

To rule out potential artifacts from ultrasound-electrode interactions during cerebellar recordings, we performed a sham ultrasound stimulation at the end of the experiment by directing the ultrasound at the upper part of the electrode array while it was still inside the brain. Our results confirm that real neural signals could be clearly distinguished from the effect of ultrasound-electrode interactions (Supplementary Fig. 2).

### 2. Different PRF of cerebellum-targeted tFUS can elicit different hippocampal neural response

To further investigate the tFUS neuromodulation effect on hippocampus from cerebellum, we recorded 62 hippocampal neurons from 13 rats subject to PRF of 1kHz (LFUS) and 10kHz (HFUS), respectively, in cerebellum-targeted tFUS (Fig. 3A). The recording loci were confirmed in the coronal brain section after H&E staining (Fig. 3B).

**Fig. 3.**
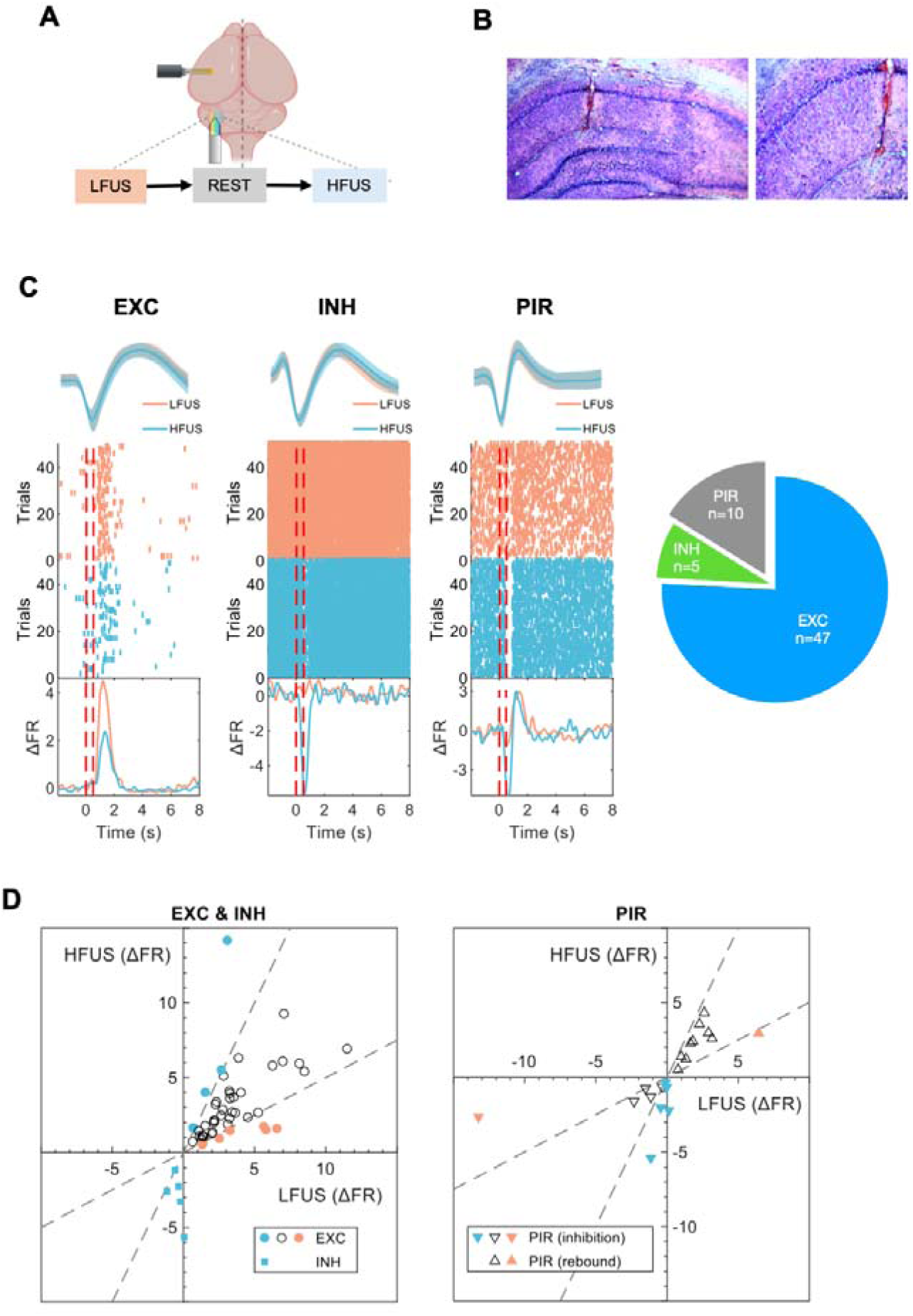
Effect of PRFs on hippocampal neural response. (A) Hippocampal neural recordings were performed with two different PRFs for tFUS at cerebellum (LFUS: 1kHz and HFUS: 10kHz) (N=13 rats). (B) Location of electrode placement shown by H&E staining. (C) Three types of hippocampal neural response (excitatory: EXC, inhibitory: INH and post-inhibitory rebound: PIR) upon cerebellum-targeted tFUS. A total of 62 units were recorded (EXC: 47, INH: 5 and PIR: 10). (D) Comparison of change in firing rate (ΔFR) of each type of hippocampal neurons upon LFUS and HFUS, respectively. The dashed lines represent ΔFR_LFUS_/ΔFR_HFUS_ = 2 or ΔFR_HFUS_/ΔFR_LFUS_ = 2. We considered a ratio of ΔFR smaller than 2 as no significant preference to LFUS or HFUS. The neurons with different PRF preference for each type are labeled with different colors in the plot.

Our experiments revealed three types of response in the hippocampal neurons (Fig.3C, Supplementary Fig.3). The first type showed increased in firing rate which started at around 0.5sec after the onset of tFUS and lasted for up to ∼1sec (Type EXC). The second type showed initial suppression of neural spikes and followed by a post-inhibitory rebound (PIR) in the firing rate (Type PIR). The third type showed suppression without PIR (Type INH). Type EXC represents the majority of the responsive neurons in our results (47/62).

Interestingly, our results showed that the magnitude of neural response could be altered by the PRF of tFUS. Among the EXC neurons (Fig. 3D, left panel), the response ratio 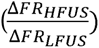 was indeed highly dispersed, ranging from 0.25 to 4.59. The median ratio of the response was 0.93, showing a weak preference to LFUS. In fact, for majority of the EXC neurons (36/47), the response ratio was within two-fold (the region between the two dashed lines in Fig. 3D). For the INH neurons (Fig. 3D, left panel), all of them showed strong preference to HFUS 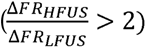. In fact, LFUS resulted in minimal response in most of these neurons. On the other hand, PIR neurons showed stronger initial inhibition to HFUS 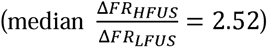. Fig. 3D (right panel) shows that in 5 out of 10 PIR neurons, the initial inhibition upon LFUS was lower than half of that upon HFUS in magnitude. However, no clear PRF preference was observed in the subsequent post-inhibitory rebound phase. The ratio of response of the PIR phase generally remained within two-fold. Overall, our results suggest that HFUS (PRF = 10kHz) at cerebellum favored inhibitory response in hippocampus.

### 3. Cerebellar cortex and DCN have different contribution to the neuromodulation effect at different PRF

To capture how different cerebellar regions may contribute to the tFUS mediated neural response in the remote hippocampal area, we injected GsMTx4 in the cerebellar cortex or the deep cerebellar nucleus (DCN) in two groups of rats respectively to block the mechanosensitive ion channels (Fig. 4A, Supplementary Fig.4). We then compared the hippocampus neural response before and after GsMTx4 microinjection in these rats.

**Fig. 4.**
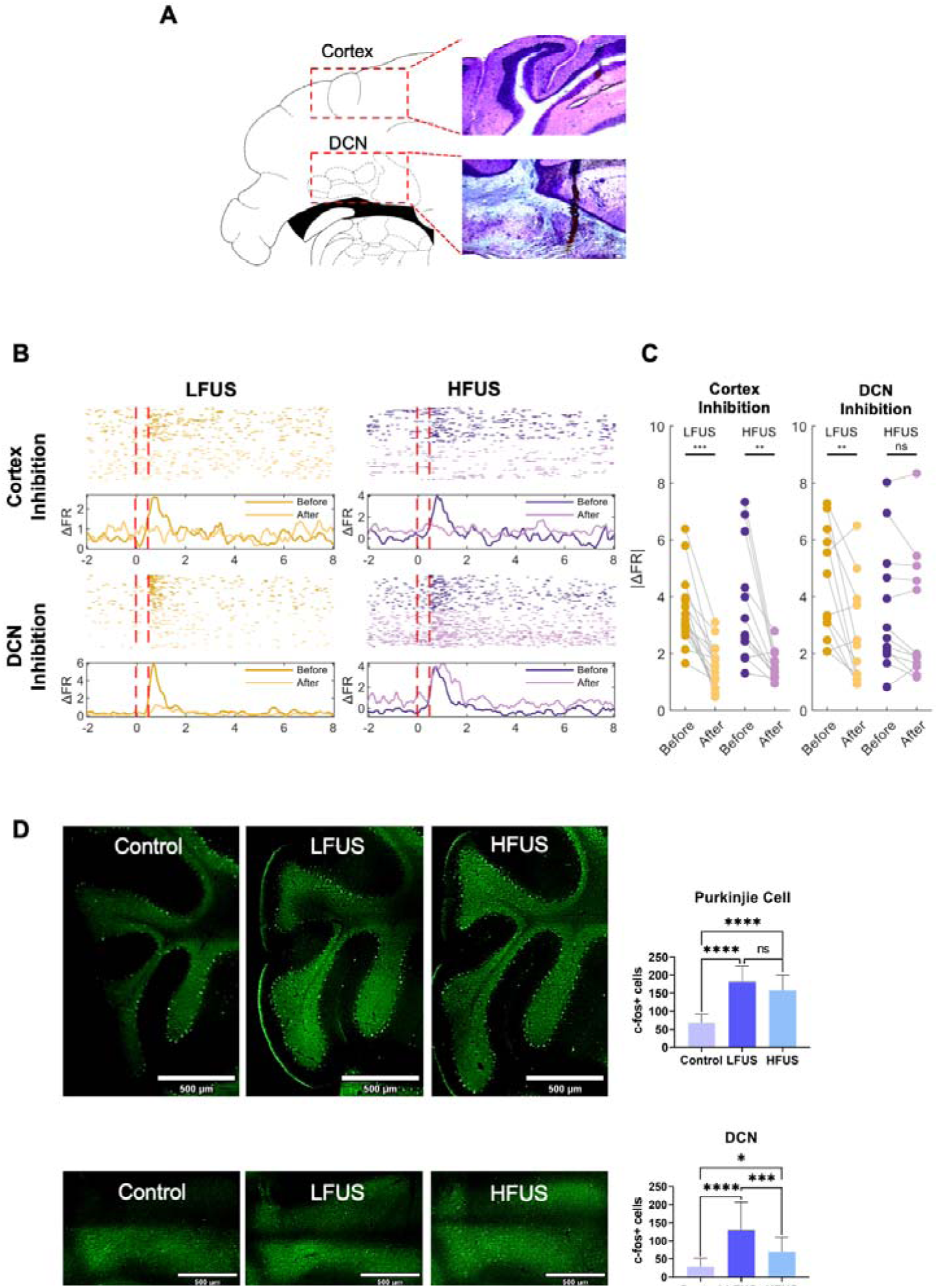
Effects of GsMTx4 microinjection in cerebellar cortex and deep cerebellar nucleus on the hippocampal neural response. (A) Loci of GsMTx4 microinjection (N=3 rats per group) by H&E staining. (B) Responses of representative hippocampal neurons following GsMTx4 inhibition in the cerebellar cortex or DCN subject to cerebellum LFUS and HFUS. (C) Summary of GsMTx4 effects on hippocampal neuron responses across all recorded units (for cerebellar cortex inhibition, n=15; for DCN inhibition, n=12). All three types of units (EXC, INH and PIR) are pooled together in the analysis. (paired t-test, *p<0.05, **p<0.01, ***p<0.001). (D) Comparison of cFos expression in Purkinje cells and DCN after LFUS or HFUS. (*p<0.05, **p<0.01, ***p<0.001, ****p < 0.0001)

Fig. 4B shows the response of one sample hippocampal neuron from each group of the rats subject to cerebellar LFUS and HFUS, respectively. When GsMTx4 was injected in the cerebellar cortex, the response in the hippocampal neuron was almost completely suppressed, regardless of the PRF used in the tFUS. On the other hand, when the DCN region was inhibited, the neural response upon LFUS was suppressed but the response upon HFUS was retained. Fig. 4C summarized the effects of GsMTx4 at cerebellum on all the hippocampal neurons that we have recorded. We have recorded both excitatory and inhibitory types of neurons in these experiments. The overall trend is consistent with the sample neurons presented in Fig. 4B. The results further confirmed that cerebellar cortex inhibition led to weakened hippocampal neural response at both PRF (n=15; LFUS: p<<0.001; HFUS: p=0.0036). On the other hand, when DCN was inhibited, the neural response was consistently suppressed upon LFUS (n=12; p=0.0065) but not HFUS (n=12; p=0.0924). These results suggested that the ultrasound mediated activation in the cerebellar cortex is more critical than in the DCN for the neuromodulation of hippocampus at high PRF of 10kHz.

We then analyzed the cFos expression in the cerebellar cortex and DCN following 20 min of tFUS to examine the spatial distribution of activated neurons in the targeted area. From 3D imaging of the cleared whole brain samples, we observed prominent cFos expression in the cerebellar cortex (Fig. 4D, upper panel) and DCN (Fig. 4D, lower panel). When comparing with the control brains, cFos expression was significantly higher in the Purkinje cells of the stimulated brains for both PRFs, showing effective activation by the tFUS on these neurons. But there was no significant difference observed between the two PRF conditions. We also observed significant cFos expression in DCN after tFUS, however, cFos expression was higher in LFUS brains than HFUS (n=3; p=0.0007), which could be partly because DCN is a deeper region.

### 4. Cerebellum-targeted tFUS activated multiple brain regions

To characterize the neural activation across different regions in the hippocampus, we analyzed the cFos expression following 20 min of cerebellum-targeted tFUS. Fig. 5A and Supplementary Fig.6 show that the cFos expression in CA1 was significantly higher after tFUS. On the contrary, it was significantly reduced in the DG. This may imply that the interneurons in DG were preferentially activated than the granule cells. No statistical difference was observed between LFUS and HFUS in either region. We did not observe significantly different expression between the ipsilateral and contralateral sides either, in both cerebellum and hippocampus (Supplementary Fig. 5). It is possibly because the ultrasound energy was not completely confined to the ipsilateral side with the Ø2mm collimator (Fig. 1C).

**Fig. 5.**
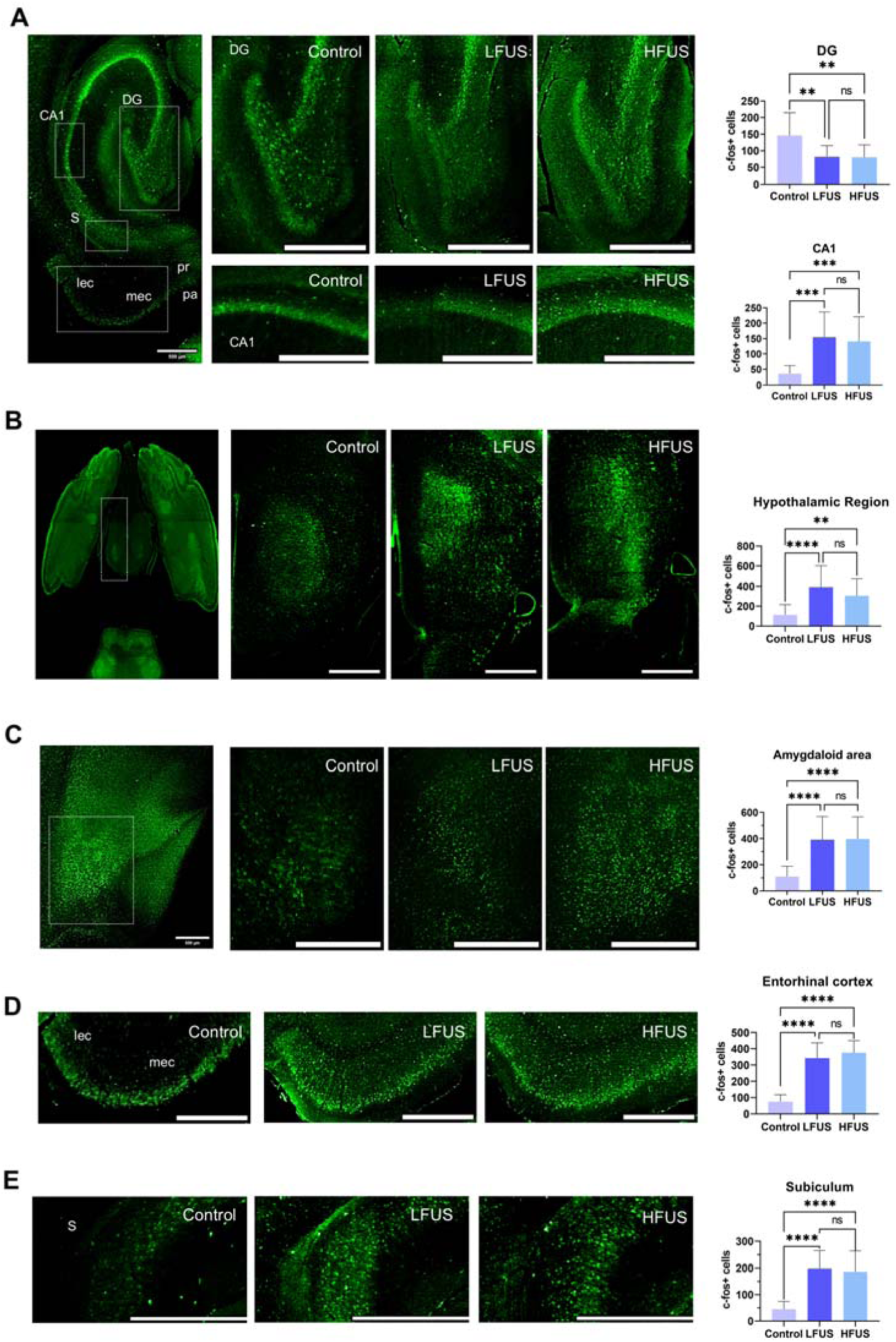
cFos expression in different brain regions after cerebellum-targeted tFUS. (A) Hippocampus (Top: Dentate Gyrus (DG); Bottom: CA1); (B) Hypothalamic region; (C) Amygdaloid area; (D) Entorhinal Cortex; (E) Subiculum. Control rats without subject to tFUS. (*p<0.05, **p<0.01, ***p<0.001, ****p < 0.0001). For whole brain cFos staining, N = 1 rat per group. For brain slice cFos staining, N = 5 rats per group.

Previous studies have shown that the cerebellum indirectly projects to hippocampus via multiple neural pathways. Here, we performed immunohistochemical analysis to assess their functional relevance in tFUS. We used 3D whole brain imaging to guide our search for brain regions that showed significant difference in cFos expression from the control group. Then we counted the number of cFos positive cells (co-stained with DAPI) from the 2D brain slice imaging in these brain regions (Supplementary Fig. 7-9). We observed that the cFos expression has significantly increased after tFUS in several brain regions. Nevertheless, we did not notice any statistical difference between LFUS and HFUS in these regions with further quantitative analysis. We found significantly higher cFos expression in the hypothalamus (Fig. 5B) and amygdala (Fig. 5C) after tFUS at cerebellum. Previous studies have revealed bidirectional connection between cerebellum and hypothalamus. In particular, it was shown that cerebellar nuclei connect to the dorsomedial and posterior hypothalamic nuclei (Cavdar, San et al. 2001). On the other hand, it was found that the cerebellum affects the activities of amygdala via a Purkinje cell-to-parabrachial nucleus pathway (Chen, Newman et al. 2023). The activated amygdala may further excite the CA1 neurons (Fig. 5A, lower part). We also observed heightened cFos expression in entorhinal cortex (EC) (Fig. 5D) and subiculum (Fig. 5E). The increased cFos in EC may follow that in the amygdala via their reciprocal connections. The EC and hippocampus have extensive reciprocal connections. The higher activity in EC may, on the other hand, be secondary to the excitation in hippocampus which sends feedback to the layer V neurons in LEC (Witter, Doan et al. 2017). Layer III neurons in EC project to the subiculum, which may explain the enhanced cFos expression in subiculum in our results. We also observed higher cFos expression in the habenular nucleus (Supplementary Fig. 9). This may result from the influence of hippocampus on habenular nucleus via the septal nucleus (Stephenson-Jones, Floros et al. 2012). Unlike the other brain regions analyzed, habenular nucleus showed high cFos expression upon HFUS than LFUS.

### 5. Cerebellum-targeted tFUS modulates cerebellum-hippocampus neural dynamics

To further examine the functional effect of cerebellum-targeted tFUS on hippocampus, we performed ERSP and PAC analysis on the LFP recorded from these two regions simultaneously. Fig. 6A shows the trial-averaged LFP power in the hippocampus and cerebellum (cortex) and the corresponding power spectra upon tFUS, respectively from the ERSP analysis. The LFP power in the cerebellum increased at the onset of stimulus. It could be observed that the power peaked in the range of ∼5-15Hz. The delayed response in hippocampus confirmed the indirect neuromodulation of tFUS from cerebellum via multisynaptic connection. The latency was lower in HFUS (∼160ms) than in LFUS (∼500ms). Both LFUS and HFUS enhanced LFP power in the hippocampus over ∼2-20 Hz. Nevertheless, LFUS-enhanced LFP power peaked in the 5-12 Hz band, while for HFUS, it peaked in the 2-5 Hz band.

**Fig. 6.**
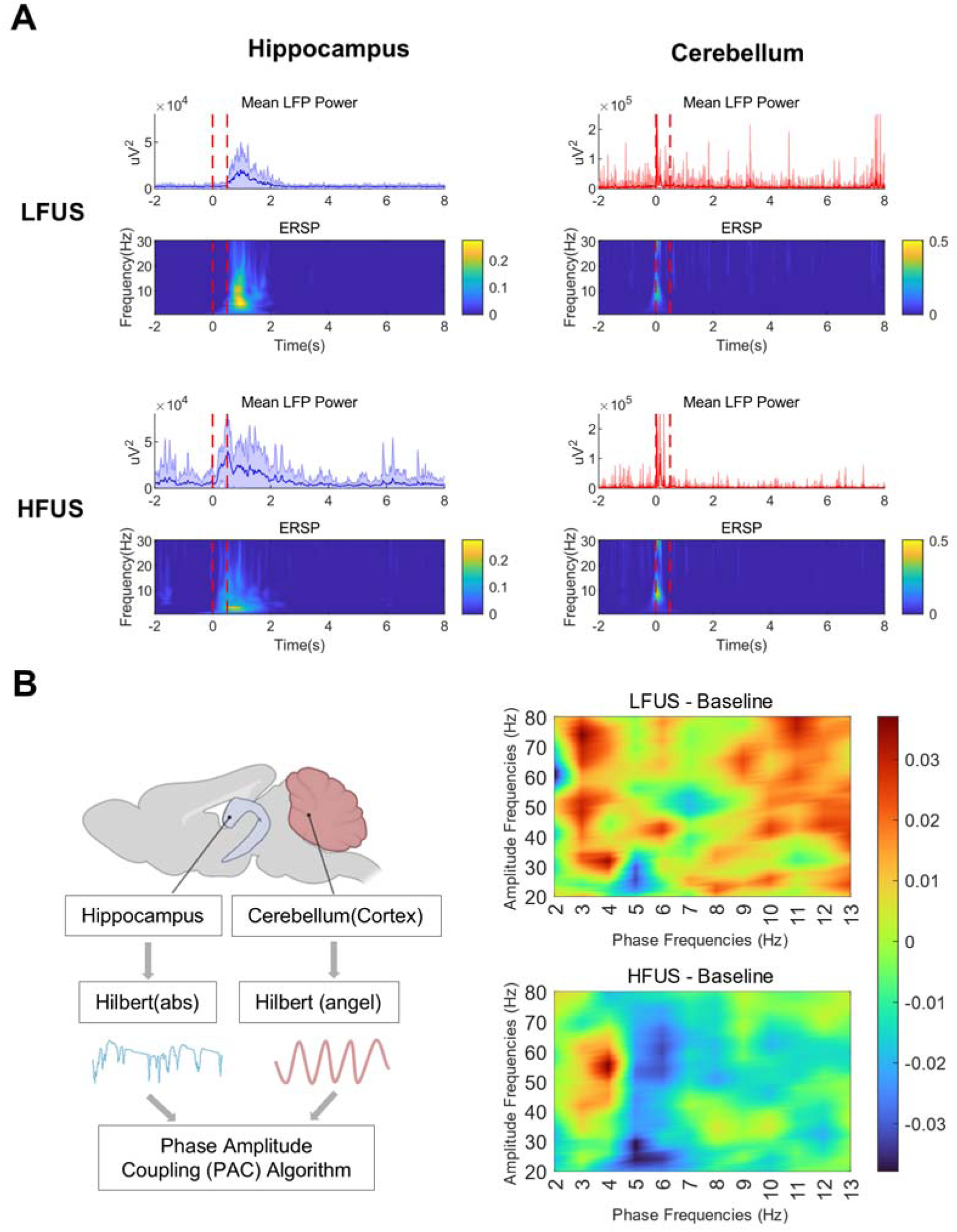
Cerebellum-targeted tFUS altered Cerebellum-hippocampus neuordynamic interaction. (A) Analysis of LFP recorded from hippocampus and cerebellum under different PRF conditions. The mean power (solid line) was computed from 60 trial of tFUS from 3 rats. The shaded area represents one standard deviation from mean. The power spectrum (bottom panel) is calculated using S-transform. (B) PAC analysis using the phase from cerebellum LFP and the amplitude from hippocampus LFP. The heatmap is plotted based on the difference in PLV between the stimulated LFP and baseline.

To further characterize the neural interaction between cerebellum and hippocampus, we performed PAC analysis between the phase of LFP in cerebellum and the amplitude of LFP in hippocampus (Fig. 6B). The heatmap shows the difference of PLVs between tFUS and baseline upon LFUS and HFUS, respectively. Under LFUS, enhanced coupling was observed over broad frequency ranges. Nevertheless, delta(1-3Hz)/alpha(8-13Hz) to gamma coupling was relatively more pronounced. On the other hand, under HFUS, delta-to-gamma coupling dominated. Furthermore, we could also observe a significant decrease in theta(4-8Hz)-to-gamma coupling under both PRF conditions. Our PAC analysis underscores the effects of different PRF in tFUS on cerebellum-hippocampus neural interactions as revealed in the different hippocampal neural responses shown in Fig. 3.

## Discussion

Our results demonstrated that cerebellar-targeted tFUS can effectively induce a variety of neural response in hippocampus. Interestingly, the cerebellum (both the cortex and DCN) was effectively activated by tFUS directly at both PRF = 1kHz and 10kHz (Fig. 4). To our knowledge, no previous report was found with effective *in vivo* neuromodulation using PRF larger than 3kHz (Tufail, Yoshihiro et al. 2011). Furthermore, cerebellum-targeted tFUS can robustly induce post-inhibitory rebound in the hippocampal neurons (Fig. 3C), which was not reported by using optical stimulation (Zeidler, Hoffmann et al. 2020). Furthermore, inhibition in individual hippocampal neurons is favored by PRF = 10kHz than 1kHz (Fig. 3D). We also found that beta-gamma coupling in cerebellum-hippocampus interaction is weakened by increasing the PRF from 1kHz to 10kHz. These results suggest that the neuromodulation effect at remote brain regions can be altered by controlling the PRF.

Previous studies have shown that neurons could respond to different PRFs of direct ultrasound stimulation. For example, Yu et al. (Yu, Niu et al. 2021) showed that excitatory neurons in the primary somatosensory cortex in mice were preferentially excited by PRF = 300Hz than 30Hz, while the opposite was observed in inhibitory neurons. On the other hand, Sherman et al. (Sherman, Bortz et al. 2024) showed that PRF selectivity was observed in individual neuron despite their types in the motor cortex of awake mice. They found that using PRFs of 10 to 140Hz, within a neuron population, most neurons responded to only one of the PRF but not the other two. The evoked calcium responses were similar between parvalbumin-positive (PV) and parvalbumin-negative (non-PV) neurons. They suggested that this is mainly attributed to the expression variation in mechanosensitive channels among individual neurons, which appears to be much more prominent than between neuron types. In another study, Murphy et al. (Murphy, Farrell et al. 2022) showed that ultrasound at PRF=900Hz increased activity of inhibitory neurons while suppressing activity of excitatory neurons in the mouse hippocampus.

Adding to these previous works, our studies first showed that tFUS at supraphysiological PRF as high as 10kHz can reliably activate neurons in rats in vivo. Interestingly, this high PRF tFUS at cerebellum favors inhibition in hippocampal neurons (Fig. 3C) and this PRF selectivity effect rely more on the cerebellar cortex than DCN (Fig. 4). Furthermore, such PRF selectivity was not due to the difference in gross activation in related brain regions (Fig. 5) but likely because of altered neural dynamic interaction (Fig. 6). Our results show that PRF selectivity of tFUS can be realized remotely. We hypothesize that we could extend the dynamic range of tFUS neuromodulation at hippocampus by using different combination of PRFs at cerebellum and directly at hippocampus, respectively. Potentially, that could achieve more robust bi-directional neuromodulation at the hippocampus, which would be useful for modulating cognitive function and for clinical application, such as epileptic management.

Our results show that the cFos expression in the DG in hippocampus decreased upon tFUS at cerebellum (Fig. 5A Upper part), while all the other brain regions, including CA1, that we have examined shown increased cFos expression. It is possibly due to accumulation of ΔFosB in DG granule cells upon repeated stimulation which suppressed further cFos reinduction (Lamothe-Molina, Franzelin et al. 2022). On the other hand, it may imply that cerebellum-targeted tFUS apparently favors activation of inhibitory interneurons in DG over excitatory granule cells, regardless of the PRF used. Previous studies showed that activating somatostatin-expressing inhibitory neurons in the hippocampus by sleep deprivation could result in reduced cFos expression in DG (Delorme, Wang et al. 2021). It has also been shown that activation of somatostatin-expressing inhibitory neurons could inhibit firing of parvalbumin-expressing interneurons in CA1 (Chamberland, Nebet et al. 2023). Nevertheless, the consequence of reduced cFos expression in DG is context-dependent. For example, the overexpression of ΔFosB could promotes resilience to stress (Nestler 2015). On the other hand, the reduced cFos may lead to persistent cognitive deficits in seizures (Corbett, You et al. 2017). Further studies will be required to examine the consequence of cerebellum-targeted tFUS on the cognitive function given this suppressed DG activities by using different animal models.

We have found that multiple brain regions were engaged by cerebellum-targeted tFUS. Nevertheless, the overall activation level, as reflected in cFos expression, did not show any PRF selectivity as in the neural spike recording. Nevertheless, our PAC analysis shows that the PRF selectivity mainly resulted in altered interaction in neural dynamics between cerebellum and hippocampus. We found that delta-gamma coupling was consistently strengthened after tFUS with both PRFs, but PRF = 1kHz also strengthened alpha-gamma coupling. Delta-gamma coupling in EEG was shown to be enhanced during the post-ictal period in seizure (Grigorovsky, Jacobs et al. 2020). However, it has not been well-studied in hippocampus specifically. On the other hand, experimental evidence suggested that alpha-gamma coupling in EEG may be associated with neuroplasticity induced by rhythmic transcranial magnetic stimulation at PRF of ∼10Hz (Wagner, Makeig et al. 2019). These couplings may suggest promising therapeutic potential for cerebellum-targeted tFUS. Nevertheless, the lower theta-gamma coupling generally implies compromised neural communication. Further studies will investigate the impact on cognitive function with these tFUS parameters.

### Limitations of the study

While our studies showed that hippocampus neurons demonstrated different response subsequent to the tFUS at cerebellum, the cell types of these neurons were not identified. We made logical speculation based on the baseline firing that the inhibitory neurons were suppressed, with or without PIR in our experiments. More sophisticated genetic engineering tools will be required to label specific cell types to distinguish, for example, somatostatin-expressing neurons from the parvalbumin-expressing interneurons.

We showed that PRF much higher than typical physiological relevant level can activate the cerebellum. However, the detailed mechanism remains unclear. Revealing the type of cells (both neurons and non-neurons) and ion channels, for example, that are responsible to engage the brain regions to such high-frequency stimulus could enrich our understanding on optimizing these transcranial stimulation tools.

## Methodology

### EXPERIMENTAL MODEL AND SUBJECT DETAILS

#### Animals

Adult Sprague Dawley rats, weighing 400g - 500 g, were used in this study. The rats were kept in standard cages in a 12-hour light/dark cycle environment and allowed free access to food and water. All procedures were performed under the approval of the Animal Research Ethics Sub-Committee of our institute. All experiments were performed in accordance with the principles outlined in the National Institutes of Health Guide for the Care and Use of Laboratory Animals.

## METHOD DETAILS

### Surgical procedure

The rats were first anesthetized with 3% isoflurane for 5 min in the isoflurane box. Then, the rat hear was fixed on a stereotaxic apparatus (Model 942, David Kopf Instruments, Tujunga, California, USA). A gas mask was attached to the rat’s nose to provide continuous isoflurane flow throughout the experiment. One or more holes were opened on the cranial for electrode implants and/or microinjection.

Connect the tube that outputs isoflurane and oxygen to the rat ‘s nose, maintained under anesthesia by1.5-3% isoflurane with ear bars and a clamping device to keep their heads horizontal during surgery. Add a 37°C heating pad under the rat’s body during the whole experiment. The fur covering each rat’s skull will be trimmed to expose the cleaner scalp, and the scalp will be sterilized with iodine. The bregma and lambda will be explored after cutting the scalp and removing the subcutaneous tissue. Cranial windows (1–2 mm in diameter) were opened in the skull using a high-speed micro drill under stereotaxic surgery assisted with microscope. One hole will be drilled at the right side of the skull as the ground. Reduce the concentration of isoflurane to 1.3%–1.5% when recording signals. Recordings were obtained using the Cerebus DAQ system (30 kHz sampling frequency, Blackrock Microsystem, USA).

### Focused ultrasound stimulation (FUS)

The fundamental frequency (FF) of the ultrasound stimulus was set by a function generator (AFG 2021-SC, Tekronix, USA). In this study, the FF was kept at 500 kHz, which is within the optimal range for stimulating intact brain (Tufail, Yoshihiro et al. 2011). The trigger signal for the stimulus was generated by a Programmable Pulse Stimulator (Master-9) which controlled the pulse repetition frequency (PRF), number of tones burst (NTB), duty cycle (DC) and interstimulus interval (ISI). The pulse signal was fed into a linear power amplifier (E&I240L, ENI Inc., USA) and transmitted to a focused ultrasound transducer (V301-SU, Olympus, USA).

A custom-designed 3D-printed collimator (diameter = 2mm) was used to focus the ultrasound wave. The ultrasound transducer was mounted on one arm of the stereotaxic apparatus above the cerebellum at an angle of 40°. Coupling gel was applied between the transducer and the cranial to maximize power transmission.

Before the experiments, a hydrophone-based (HCL-0200, Onda, USA) three-dimensional (3D) ultrasound pressure scanning system (AG-2010, Onda, USA) was used to experimentally measure the FUS spatial profile in our setup. We included the rat cranial skull but not the base of the skull in our scanning.

### Electrophysiology

A 16-channel (4-by-4) 8-mm single shank electrodes (KD-MEA-16, KEDOU, China) was used in our neural recording. The electrodes were implanted at cerebellum (ML: −4 mm, AP: −10 mm, DV: −3 mm) and/or hippocampus (ML: −2.3 mm, AP: −3 mm, DV: −3 mm) using a stereotaxic device. The broadband neural signals were acquired by the Cerebus system (Blackrock Microsystems, USA) sampled at 30 kHz. The neural spikes were obtained by highpass filtering at 350Hz, and the the local field potentials (LFP) were obtained by lowpass filtering at 250Hz and downsampled to 1kHz. Spike sorting was performed using the Blackrock Offline Spike Sorter (BOSS) software. Time stamped event flags from the Programmable Pulse Stimulator (Master-9) related to the tFUS were also recorded simultaneously with the neural signals.

### Histology

About 90min after tFUS, the rats were deeply anesthetized with an overdose of anesthetics and transcranial perfused with phosphate buffer saline (PBS, 0.01 M, pH 7.4) and 4% paraformaldehyde (PFA, Thermo Fisher Scientific, Cat. #119690010). The brains were then carefully extracted and post-fixed in 4% PFA for 24h, dehydrated in 30% sucrose for 48h. The fixed brains were cut into 25μm slices with a freezing microtome for H&E, Nissl, and immunofluorescence staining.

### Microinjection of GsMTx4

We used GsMTx4 (Cat. No.: HY-P1410, MCE), a selective inhibitor of cationic mechanosensitive channels including TRP and Piezo families, to inactivate the cerebellum and/or hippocampus from the ultrasound stimulation. GsMTx4 powder 250μM. Microinjections of the GsMTx4 were performed using glass pipettes attached was dissolved in phosphate buffer saline (PBS 1×) to prepare a GsMTx4 solution at to a 5µL Hamilton microsyringe under the control of a stereotaxic microinjection pump. After each injection (0.5 μL), the pipette was left in place for 10 mins before it is withdrawn from the brain.

### Brain slice cFos immunofluorescence staining and imaging

To minimize nonspecific antibody binding, brain slices were first blocked in blocking buffer (10% normal goat serum, 5% BSA, and 0.3% Triton 1×PBS) for 2h at room temperature (RT). After PBS washes (3×10 min), sections were incubated with cFos primary antibody (1:1000; Cell Signaling Technology, Cat. #2250) diluted in blocking buffer for 16–20h at 4°C. Following PBS washes, sections were incubated with Alexa Fluor 555 goat anti-rabbit IgG (H+L) secondary antibody (1:1000; Invitrogen, Cat. #A21428) diluted in blocking buffer for 2h at RT with shaking. After PBS washes (3×5 min), nuclei were stained with DAPI (Invitrogen, Cat. #R37606). Sections were mounted with antifade fluorescence mounting medium (Abcam, Cat. #ab104135) and stored at 4°C until imaging within two weeks.

All coronal brain sections were imaged using a Leica TCS SP8 Confocal Microscope. Using a 10× objective, a series of overlapping images of the entire coronal section were acquired and stitched into a single composite image.

### Whole brain cFos immunofluorescence staining and imaging

Whole rat brains were immunostained following the INSIHGT workflow (Yau, Hung et al. 2024). The brains were fixed in 4% PFA at 4°C for 24h, then washed in PBSN (0.02% w/v sodium azide in 1× PBS) at RT with gentle shaking. Brain dehydration was performed sequentially in 50% methanol for 30 min, followed by three 30-min washes in 100% methanol at RT with shaking. Delipidation was performed by incubating brains in a 2:1 (v/v) mixture of dichloromethane/methanol at RT for 24h. Subsequently, brains were rehydrated through reversed dehydration steps and washed with PBSN.

Following overnight incubation in 1× INSIHGT buffer A at 37°C with shaking, the brain samples were simultaneously incubated with anti-cFos primary antibody (1:1000; Cell Signaling Technology, Cat. #2250) and AffiniPure Fab Fragment Donkey Anti-Rabbit IgG (H+L) secondary antibody (Jackson ImmunoResearch, Cat. #711-547-003) in staining buffer for 8 days at RT under light-protected conditions. Subsequently, brains were incubated in INSIHGT buffer B overnight. The stained brains were washed in PBSN and dehydrated through graded methanol (25%, 50%, 80%, and 100%) for 8h each with continuous shaking at RT, followed by a final 12h incubation in 100% methanol. The samples were then incubated in 100% dichloromethane for 48-60h at RT.

Optical clearing was achieved by incubating brains in a 1:2 (v/v) mixture of benzyl alcohol/benzyl benzoate (BABB) for 48h until samples became fully transparent. Cleared brains were imaged using a custom-built MesoSPIM v5.1 (Voigt, Kirschenbaum et al. 2019). Images were acquired at 2× magnification, with a pixel size of 3.224 × 3.224 µm and an optical plane spacing of 2 µm.

### Neural firing rate analysis

The firing rate of the neurons was calculated from the peri-stimulus time histogram (PSTH) using a bin of 50 msec. To compare the neural response, we computed the change in firing rate from the baseline rate (averaged over 2sec before tFUS) and then normalized to the baseline rate:

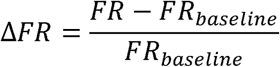

### The Event-Related Spectral Perturbation (ERSP) analysis

ERSP was used to measure the average dynamic changes in amplitude of a broadband LFP frequency spectrum as a function of time relative to a tFUS event. The LFP was first downsampled to 500Hz. Then it was notch-filtered to remove the 50Hz noise, before it was bandpass filtered within 2-40Hz. We used S-transform to provide the joint time-frequency distribution (TFD) of the LFP power. The mean baseline level was computed by averaging the power spectra over the 2sec before tFUS. Then power spectra for each trial was subtracted by the corresponding mean baseline spectra. The normalized spectra were then averaged over all the trials.

### Phase-amplitude coupling (PAC) Analysis

The PAC analysis examines how the phase of one low frequency time series signal drives the amplitude of another high frequency signal. Here, we use Phase Locking Value (PLV) to quantify the cerebellar phase-hippocampal amplitude coupling. We 20-80 Hz. The phase of the filtered cerebellar LFP 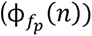 was extracted. On the first bandpass filtered the cerebellar LFP at 2-13 Hz, and the hippocampal LFP at hippocampal LFP which was then Hilbert transformed. The phase 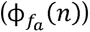 of the other hand, the amplitude time series signal was first extracted from the filtered transformed signal was then extracted. Then the PLV was computed as follow (Mormann, Fell et al. 2005):

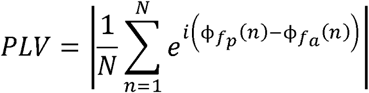

where *N* is the length of the time series signal and *n* is the time index.

### Brain slice cFos quantification

The number of cFos-positive cells in the brain slices was quantified using the QUINT workflow (Yates, Groeneboom et al. 2019) for automated cell quantification (Supplementary Fig. 7B). Preprocessing was performed using FIJI (Schindelin, Arganda-Carreras et al. 2012) to prepare the data for atlas registration and cell detection. Atlas registration was carried out by aligning individual sections to Waxholm Atlas of the Spraque Dawley Rat (WHS v4) (Kleven, Bjerke et al. 2023) using the QUICKNII software (Puchades, Csucs et al. 2019), generating customized atlas maps for each section. These atlas maps were subsequently used to analyse the distribution of cFos-positive cells across the brain. To identify cFos-positive cells, images of brain sections were segmented using the pixel classification function of ilastik software (Berg, Kutra et al. 2019). During classifier training, pixels within cFos-positive cells were manually selected for 20% of the brain sections from each animal. The algorithm then predicted high-probability pixel locations, which were manually reviewed to identify and correct misclassified positive cells. Incorrectly labeled pixels were marked as artifacts and incorporated into the next round of training. This process was repeated five times per animal. The trained pixel classifier was batch-applied to the remaining brain sections. Custom atlas maps and cFos-positive segmentations were combined using Nutil Quantifier software (Groeneboom, Yates et al. 2020), to provide cFos-positive object counts for each brain region. Regions with low-quality counts were manually reanalyzed and recounted to ensure accuracy.

### Whole brain cFos quantification

Image stitching was performed using the BigStitcher plugin (Hörl, Rojas Rusak et al. 2019) in ImageJ. Automated whole-brain atlas registration and cell detection were performed using the BrainGlobe workflow (Supplementary Fig. 7A). Atlas registration was carried out using Brainreg plugin (Niedworok, Brown et al. 2016, Tyson, Vélez-Fort et al. 2022) in napari (napari Work). Atlas data were provided by BrainGlobe Atlas API (Claudi, Petrucco et al. 2020). Cell detection was performed using Cellfinder software (Tyson, Rousseau et al. 2021), which identified regions with high cFos-positive cell densities. Regions were visualized and refined using FIJI (Schindelin, Arganda-Carreras et al. 2012).

### Statistical Analysis

Data are presented as mean values accompanied by SEM. No statistical methods were used to predetermine sample sizes. Data analysis was performed blind to the conditions of the experiments. Data was analyzed using GraphPad Prism software. Statistical comparisons were performed using one-way ANOVA followed by Tukey multiple comparison post-hoc tests, unless otherwise stated. Statistical significance was indicated as *p<0.05, **p<0.01, ***p<0.001, ****p < 0.0001, ns=not significant.

## Acknowledgments

We thank Prof. Lei Sun (The Hong Kong Polytechnic University) and Dr. Chongyun Wang (Hong Kong Centre for Cerebro-Cardiovascular Health Engineering) for their support on the ultrasound pressure scanning system.

## Author contributions

C. T. and Y. L. conceived the project. C. T. supervised the project. Y. L. designed and performed the experiments. J. T. S. H. performed the whole brain imaging. H. M. L provided access to the whole brain imaging facility and supervised on the procedure. Y. L., K. Y. C. and C. Z. analyzed the data. C. T. and Y. L. wrote the manuscript with inputs from all authors.

