## Supplemental Figure 1-9 for "Cerebellum-Targeted Transcranial Focused Ultrasound Stimulation Modulates Hippocampus Neural Activities"

Supplementary Figures 1 to 9

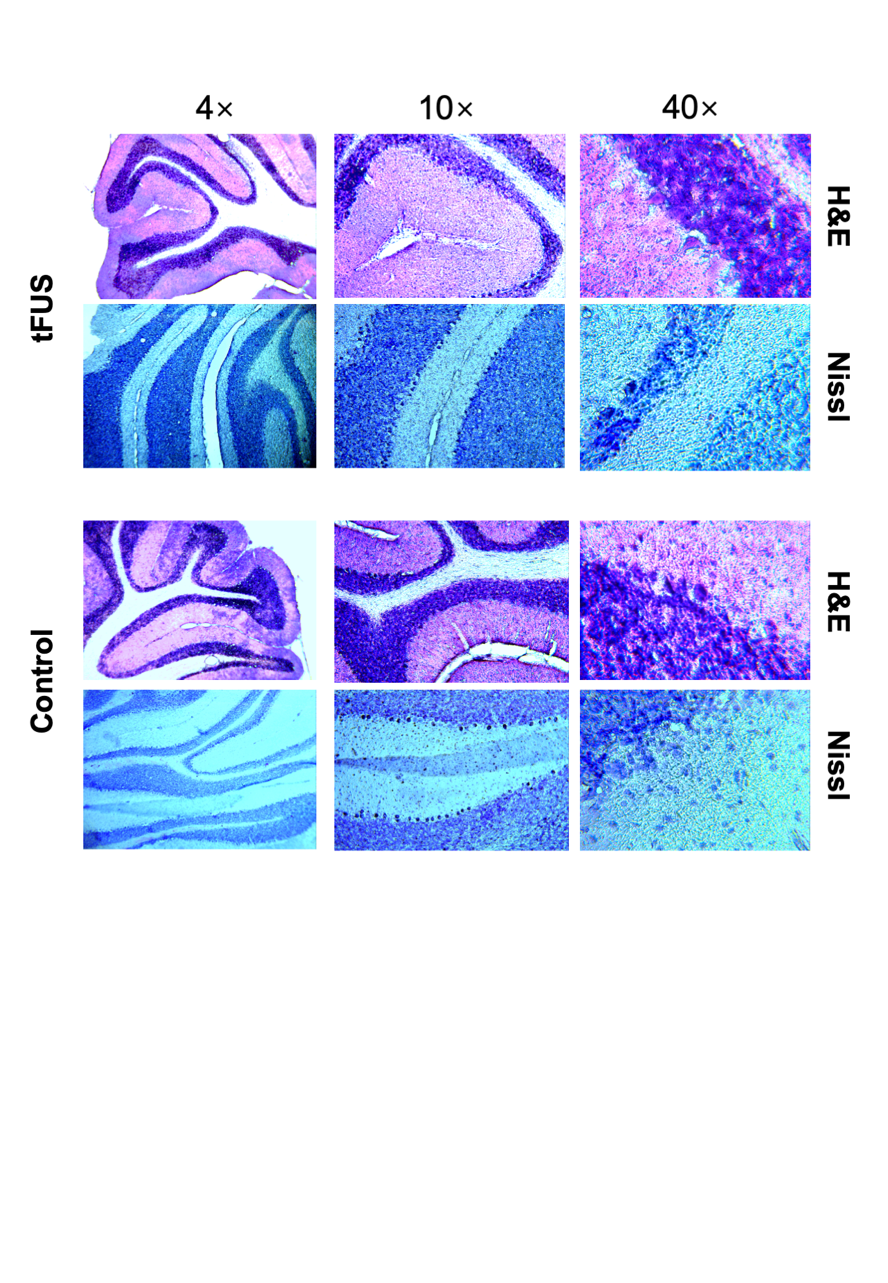

**Supplementary Fig.1** H&E/Nissl staining of cerebellum shows no tissue damage by tFUS.

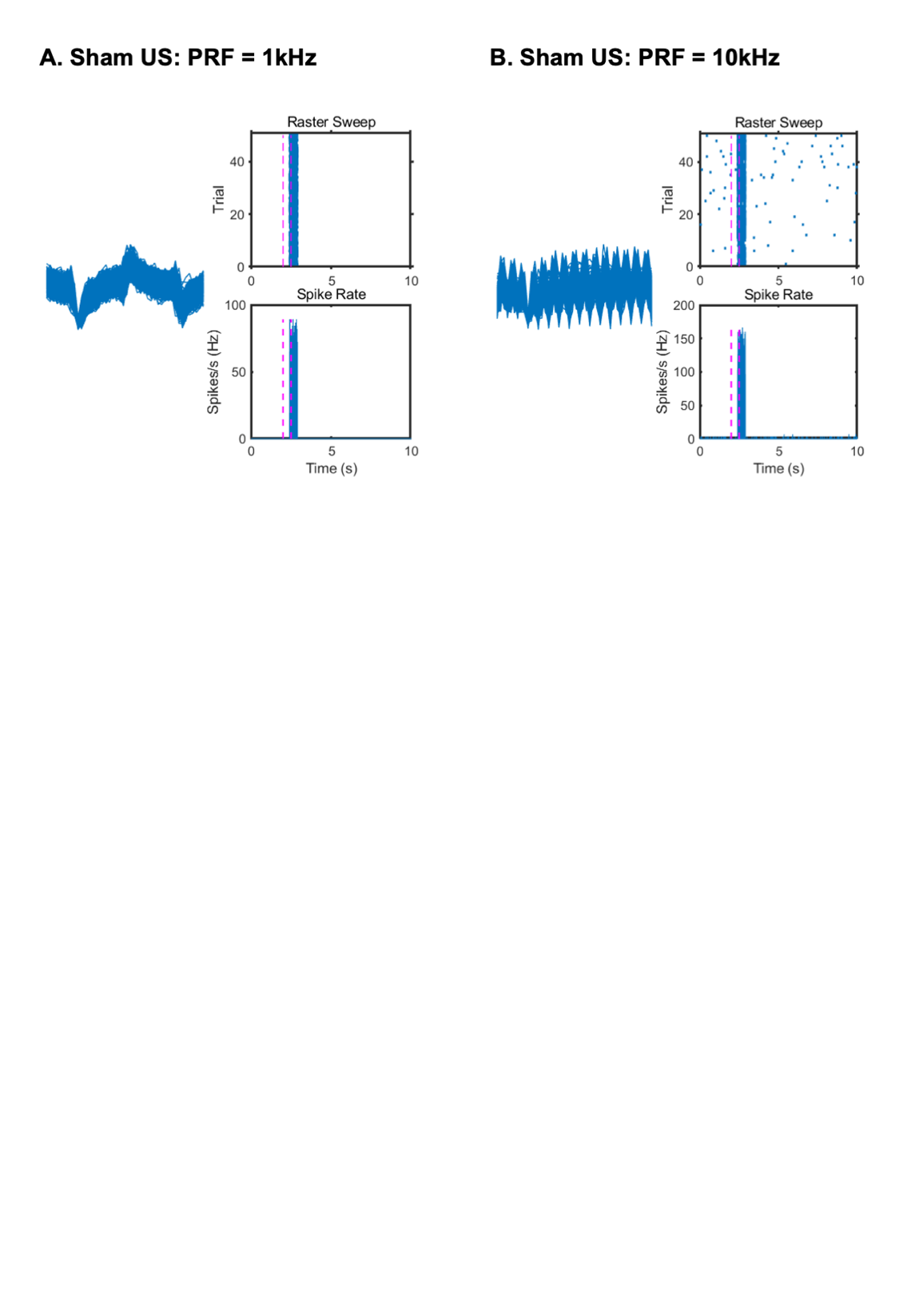

**Supplementary Fig.2** Electrode vibration-induced artifacts during sham ultrasound stimulation at two different pulse repetition frequencies (A: PRF = 1kHz; B: PRF = 10kHz).

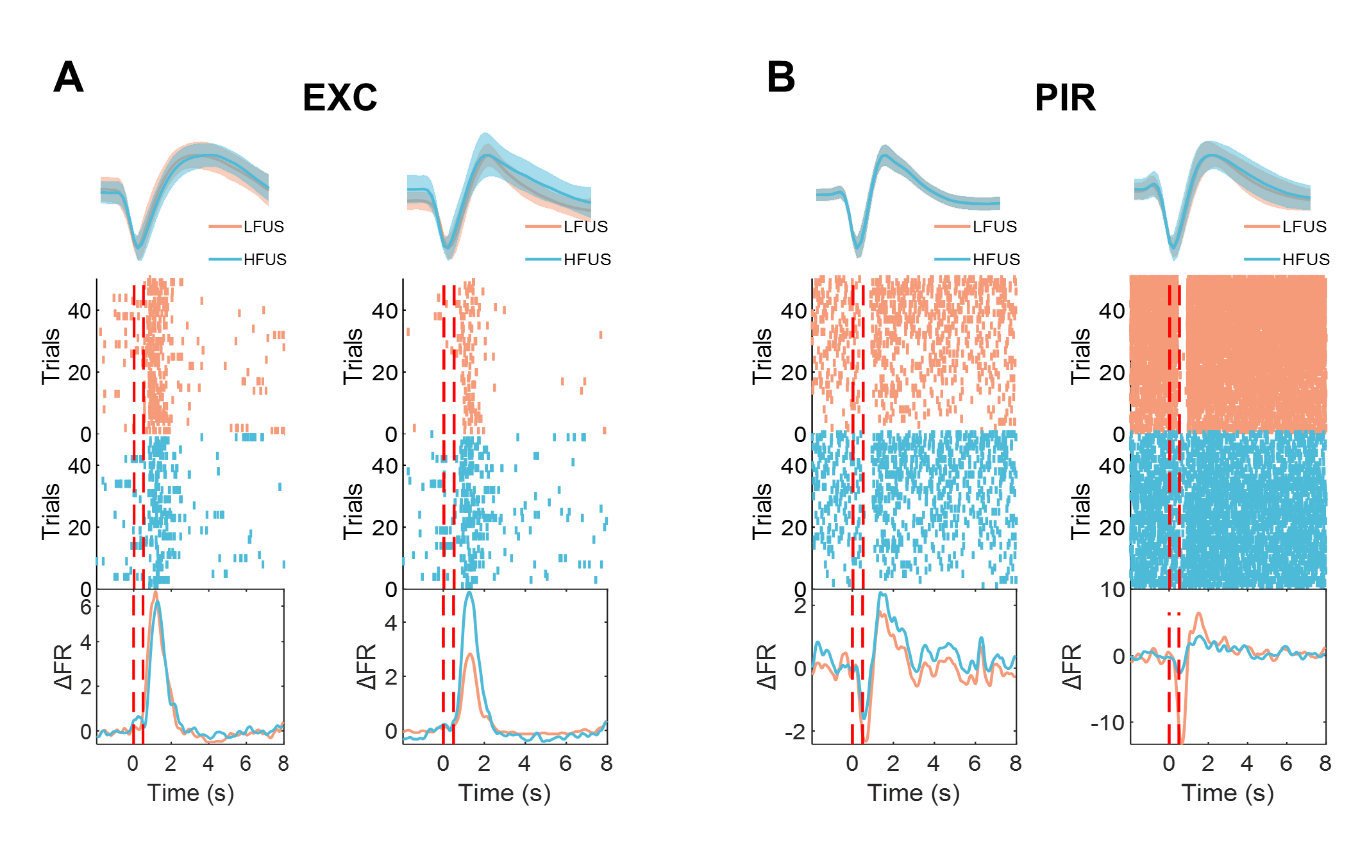

**Supplementary Fig.3 Additional representative units of type EXC and type PIR (refer to Fig. 3C).** (A) Sample EXC units showing no preference to PRF (left) and stronger response to HFUS (right); (B) Sample PIR units showing no preference to PRF (left) and stronger response to LFUS (right)

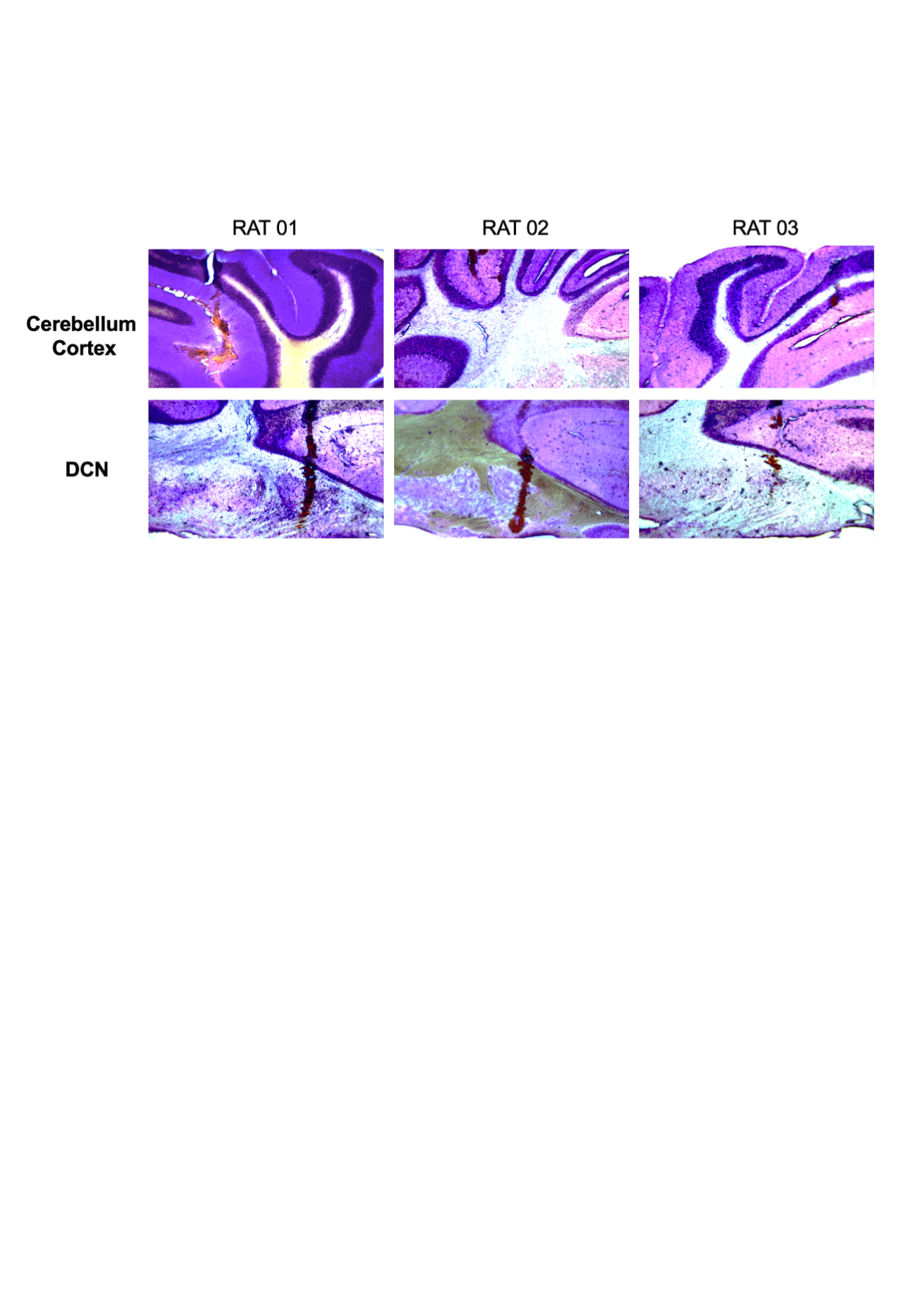

**Supplementary Fig.4** Nissl staining shows the injection site of GsMTx4 at cerebellar cortex and DCN respectively (refer to Fig. 4).

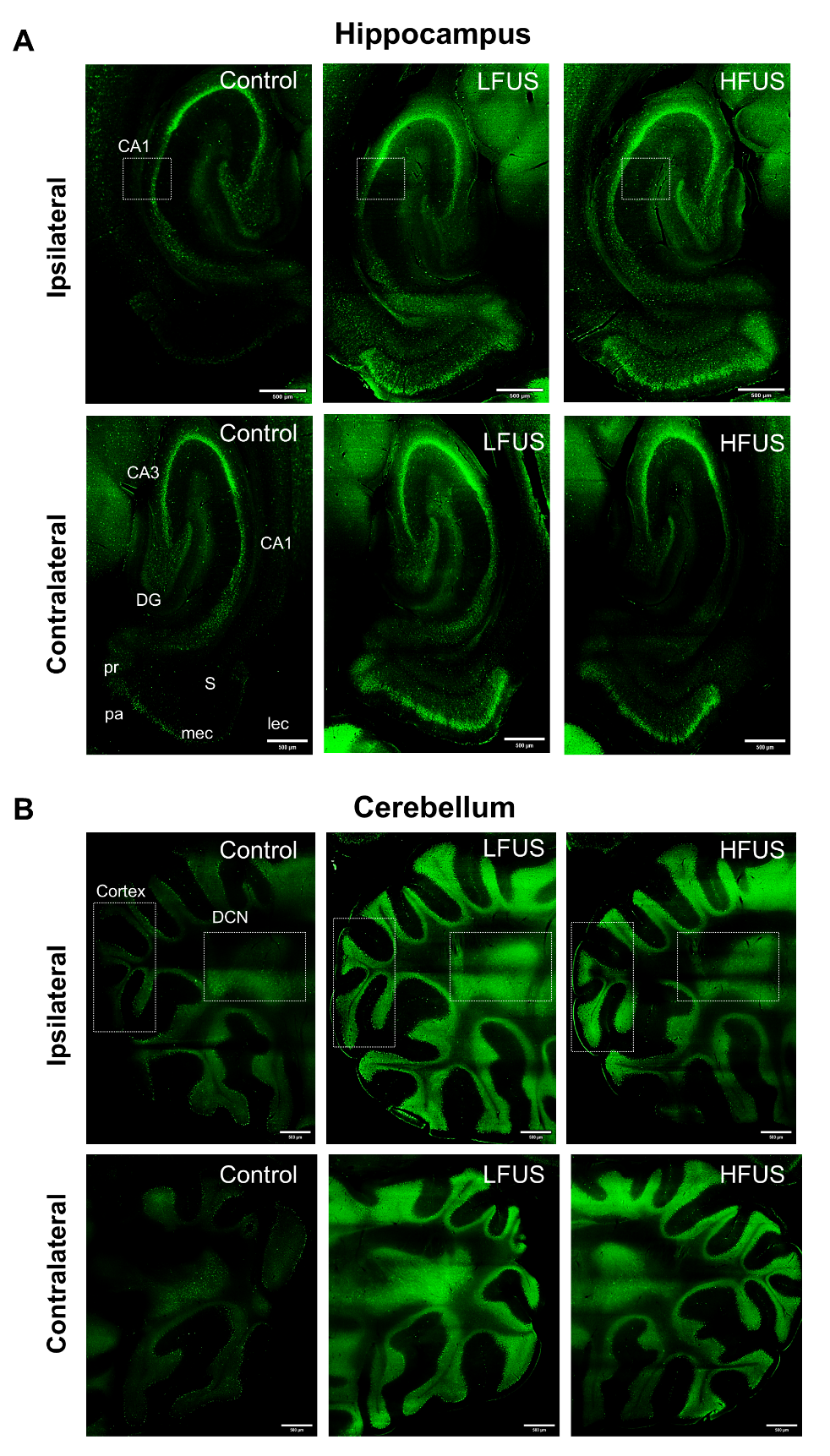

**Supplementary Fig.**5 Whole brain imaging for cFos expression in ipsilateral and contralateral hippocampus and cerebellum respectively.

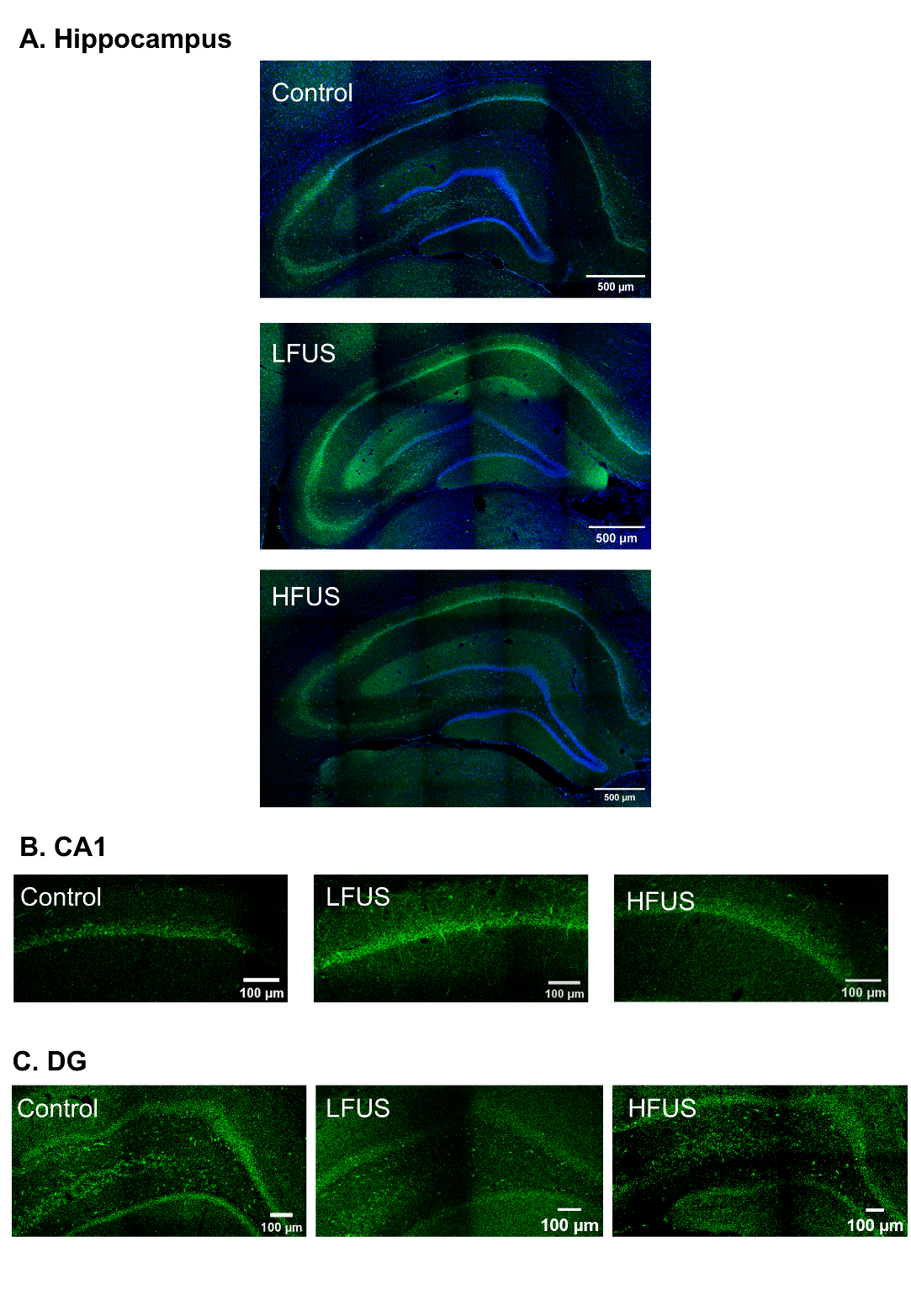

**Supplementary Fig.6** Representative brain slice imaging of cFos expression (A) Overview of hippocampus (co-stained with DAPI). (B) CA1. (C) Dentate gyrus

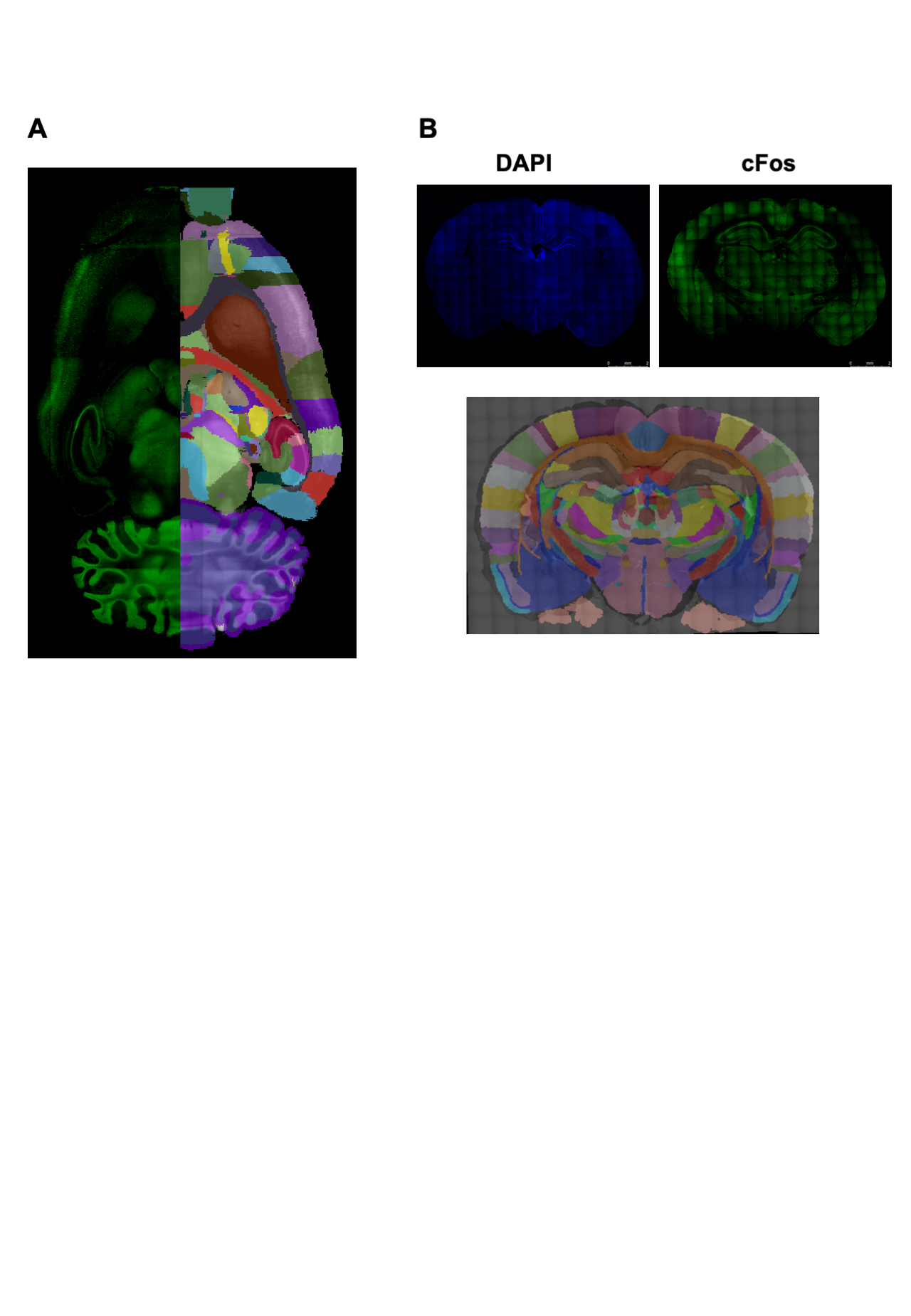

**Supplementary Fig.7 Brain imaging with precise anatomical segmentation and standardized atlas registration.** (A) 3D reconstruction of rat brain showing cFos fluorescence (left) and automated region registration (right) using BrainGlobe software. (B) Brain section analysis with DAPI nuclear staining (top left), cFos expression (top right), and automated region segmentation generated using QUICKNII (bottom).

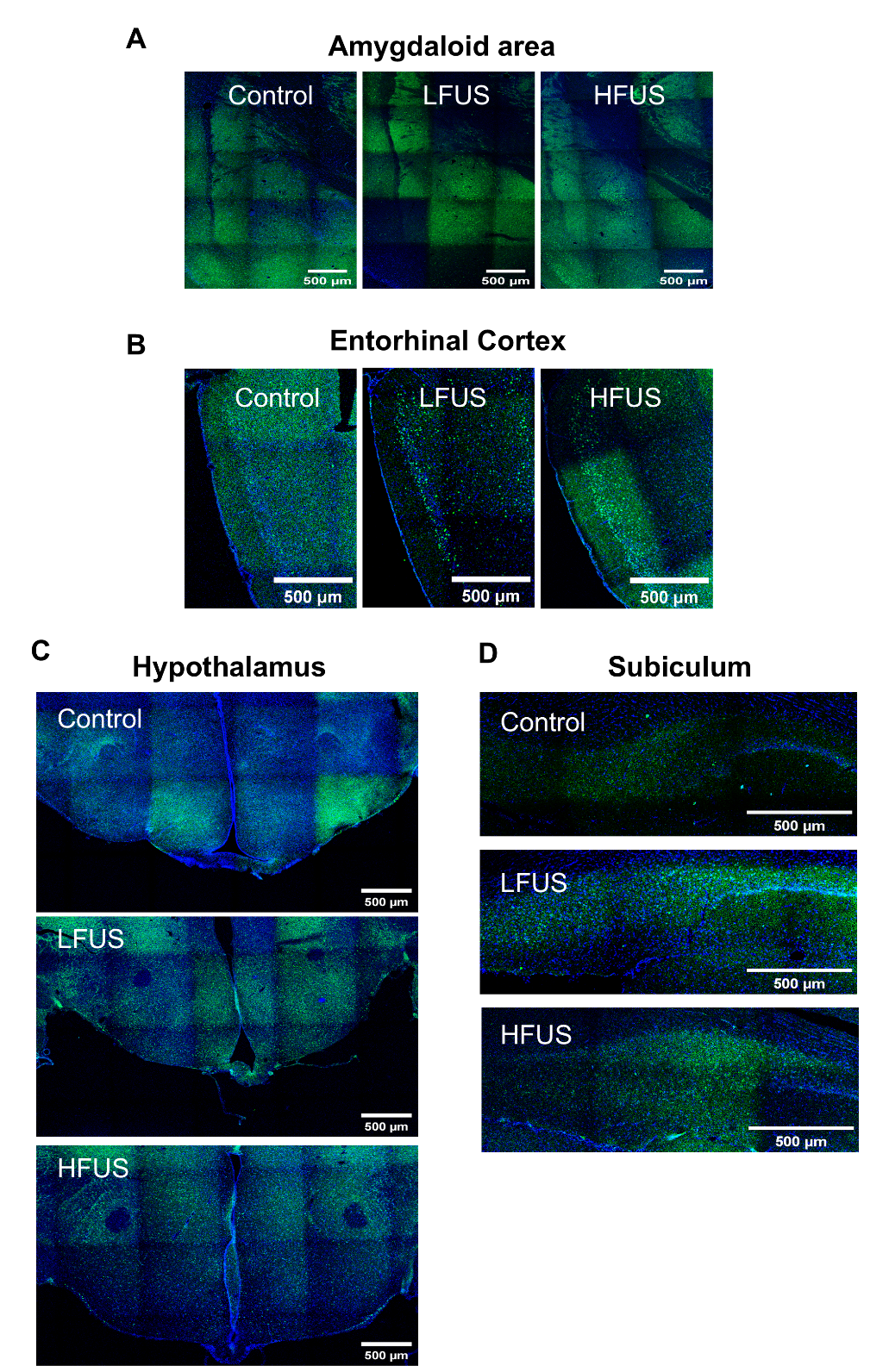

**Supplementary Fig.8** Representative brain slice imaging of cFos expression and DAPI in (A) Amygdaloid area, (B) Entorhinal Cortex, (C) Hypothalamus and (D) Subiculum (refer to Fig. 5).

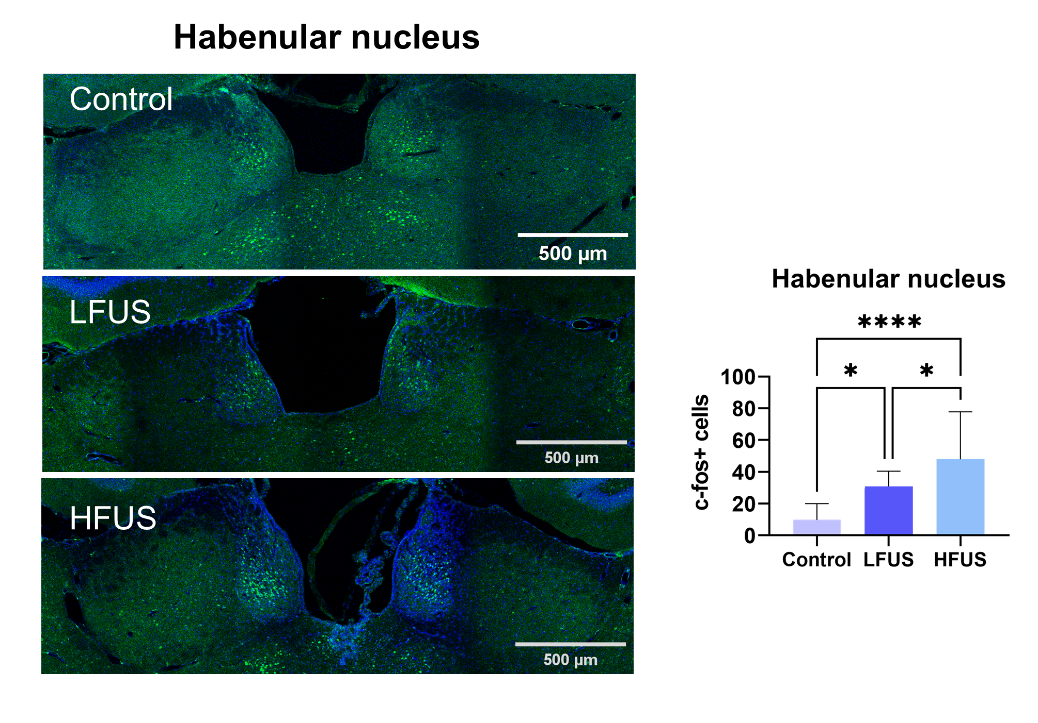

**Supplementary Fig.9** Representative brain slice imaging of cFos and DAPI in Habenular nucleus. Enhanced cFos expression in Habenular nucleus after tFUS was observed in the brain slice imaging but not whole brain imaging. (*p < 0.05, **p < 0.01, ***p < 0.001, ****p < 0.0001)
